# Mapping Kappa Opioid Receptor Binding in Titi Monkeys with ^11^C-GR103545 PET

**DOI:** 10.1101/2024.08.01.606042

**Authors:** Alita Jesal D Almeida, Brad A. Hobson, Logan E. Savidge, Claudia Manca, John P. Paulus, Karen L. Bales, Abhijit J. Chaudhari

## Abstract

**Purpose:** The kappa opioid receptor (KOR) plays a pivotal role in stress- and anxiety-related behaviors, especially in social separation and bonding. However, KOR modulations in these social contexts are not fully characterized. The coppery titi monkey (*Plecturocebus cupreus*) has been utilized as a translational animal model for studying the neurobiology of attachment, as they form socially monogamous adult pair bonds. While the PET radiotracer ^11^C-GR103545 has shown the ability to track KOR activity in other species, it has not been utilized in titi monkeys. This study assessed ^11^C-GR103545 PET for characterizing KOR activity *in vivo* and its pharmacological blockade in titi monkeys.

**Methods:** Adult titi monkeys (N=6) underwent ^11^C-GR103545 PET brain scans at baseline, followed by repeat scans upon administration of a KOR antagonist (CERC-501) and a KOR agonist (U50,488). Region-specific, non-displaceable binding potential (BP_ND_) was calculated, with the cerebellum as the reference region, focusing on 14 brain volumes of interest (VOIs) implicated in social bonding.

**Results:** Baseline ^11^C-GR103545 uptake characteristics across VOIs were consistent with previous reports in humans and other monkey models. CERC-501 pretreatment led to a significant reduction in BP_ND_ (average 55.99%) across several, but not all brain VOIs, with the most notable reduction in the superior frontal gyrus (76.25%). In contrast, U50,488 pretreatment resulted in no significant BP_ND_ changes across the VOIs analyzed.

**Conclusions:** This study demonstrates the utility of ^11^C-GR103545 to assess KOR binding dynamics in a monkey model of social bonding. Region-specific blockade of KOR was observed after pretreatment by CERC-501, while blockade by the KOR agonist U50,488 did not change ^11^C-GR103545 binding.

## Introduction

Kappa opioid receptors (KORs), the most abundant sub-type of opioid receptors, and their endogenous ligand dynorphin, play a central role in a wide range of neurophysiological functions pertaining to regulation of mood and motivation [1, 2]. KOR activation or agonism, particularly in response to experiencing stress, has been associated with aversive mood disorders [3, 4] while KOR antagonism or deactivation has demonstrated anti-depressive-like effects [5, 6]. Loneliness experienced during the COVID-19 pandemic has emphasized the amplitude of psychological distress that can be caused from the lack of human interaction [7]. Therefore, now more than ever, understanding the role of KOR perturbations in attachment relationships, preferably *in vivo*, has become crucial. While studies have assessed KOR modulations [8, 9], the underlying mechanisms remain largely elusive.

The coppery titi monkey (*Plecturocebus cupreus*), a socially monogamous South American non-human primate, exhibits several characteristics of a human adult pair bond including preference for their familiar partner, distress upon separation (social separation), and the ability of the partner to alter their stress response (social buffering) [10]. They have also demonstrated similar oxytocin release patterns to that of humans [11], making them robust animal models to analyze KOR (and oxytocin associations) in the broader context of social relationships.

Positron Emission Tomography (PET) has been used to evaluate KOR changes in studies of humans with substance abuse and major depressive disorders [12, 13], but not in the context of social attachment. Additionally, to date, no PET studies analyzing the KOR have been conducted in titi monkeys. To address this unmet need, the aims of this study were to (i) characterize the distribution and binding of ^11^C-GR103545, a KOR agonist PET radiotracer, in the brain of titi monkeys, and (ii) determine the effects on ^11^C-GR103545 binding post-pharmacological blockade of the KOR with a KOR antagonist and agonist.

## Methods

### Animals

Six adult (7.0±1.5 year old) coppery titi monkeys (3 male, 3 female), weighing 1.22±0.15 kg, housed at the California National Primate Research Center, were selected for this study (details in **Supplementary Section S.1)**.

### Imaging protocol and treatment

The PET radiotracer ^11^C-GR103545 was synthesized using the method described in [14]. PET data acquisition began approximately 15-s prior to administration of ^11^C-GR103545 with an average injected activity of 48.16±6.27 MBq. Blocking experiments were performed with KOR antagonist CERC-501 (0.3mg/kg, i.m.) and KOR agonist U50,488 (3 mg/kg, i.m.), injected 10-min and 20-min respectively, prior to radiotracer administration [15]. Imaging sessions for each animal were spaced at least one month apart. To aid with regional delineation, an anatomical T_1_-weighted brain MRI scan was acquired (details of PET and MRI acquisitions are in **Supplementary Sections S1.2-1.3**).

### Image post processing and kinetic modeling

A titi monkey brain atlas was warped to each animal’s anatomical MRI, producing regional labels. Region-specific, non-displaceable binding potential (BP_ND_) was calculated utilizing the simplified reference tissue model (SRTM) (**Supplementary Section S1.4**), with the cerebellum as the reference region. Primary analysis included 14 brain volumes of interest (VOIs) that have been implicated in social attachment, namely, nucleus accumbens, caudate, hypothalamus, cingulate, hippocampus, amygdala, claustrum, orbitofrontal cortex, thalamus, septum, superior frontal gyrus, superior temporal gyrus, insula, and putamen. Additionally, outside the brain, the pituitary gland was delineated by drawing a sphere due to the high PET signal observed in that region. Secondarily, Logan reference tissue model (LRTM) was also utilized for BP_ND_ calculation for comparison with other published data [15] (**Supplementary Section S1.5**).

### Statistical analysis

A one-way nonparametric ANOVA test for matched pairs (Friedman test) was run (GraphPad Prism v10.0.2) for BP_ND_ comparisons between baseline versus after antagonist and agonist pretreatment [16]. Multiple comparisons were accounted for by controlling the false discovery rate (FDR) [17]. The significance threshold was set at 0.05.

## Results

### Baseline BP_ND_ concurred with reported distribution of KOR in the titi monkey brain

The reported BP_ND_ patterns from baseline scans (**Figure 2**), were consistent with autoradiography studies reporting KOR density in the titi monkey brain [18]. High binding (BP_ND_ > 1.3) was observed in the pituitary gland followed by the insula, claustrum and orbitofrontal cortex. Moderate binding (0.9 < BP_ND_ < 1.3) VOIs include (but are not limited to) the nucleus accumbens, amygdala, and hippocampus. Lastly, the thalamus had low binding (BP_ND_ < 0.9).

**Fig. 1.**
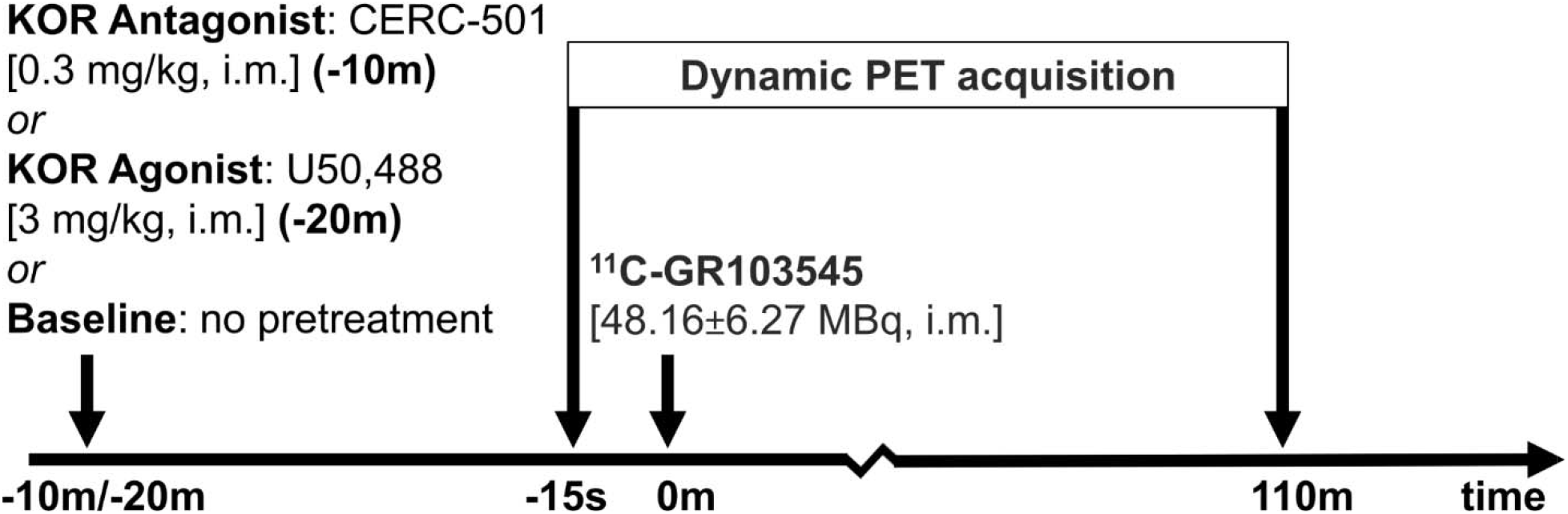
Schematic showing the scanning paradigm for N=6 titi monkeys, each imaged for a duration of 110-min (PET emission scan), at 3 timepoints; baseline, 10-min after pretreatment with antagonist CERC-501 and 20-min after pretreatment with agonist U50,488. Imaging sessions for each animal were spaced at least one month apart of each other

**Fig. 2.**
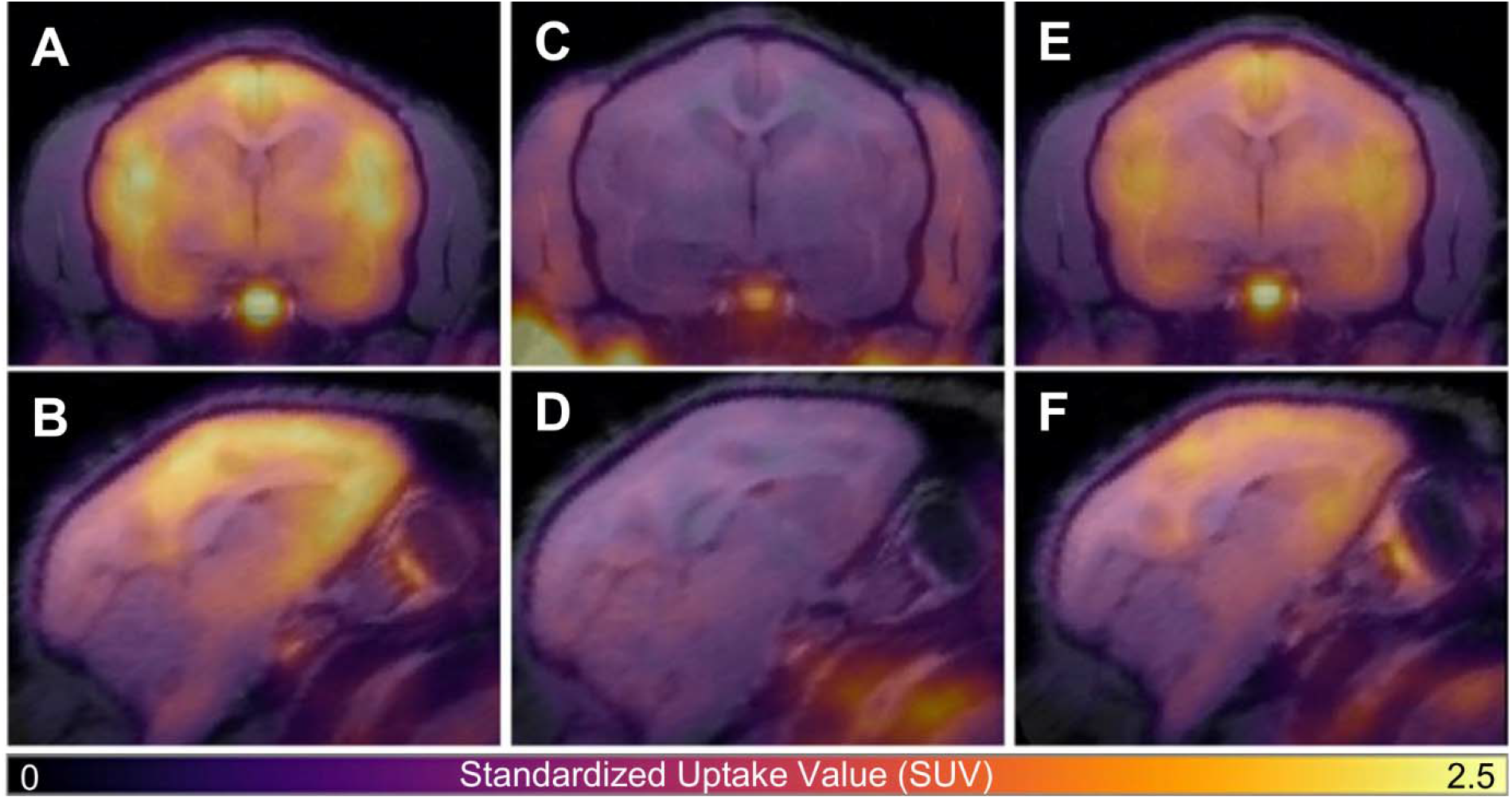
^11^C-GR103545 PET (color) overlaid onto T_1_-weighted MRI of a titi monkey in the axial (A, C, E) and sagittal (B, D, F) plane for baseline (A, B), antagonist (C, D) and agonist (E, F) pretreatment scans. Reduced radiotracer uptake in the brain is apparent after pretreatment with the KOR antagonist (C, D). PET data are presented as standardized uptake value (SUV) images calculated from data 80-110 min post-radiotracer injection

**Fig. 3.**
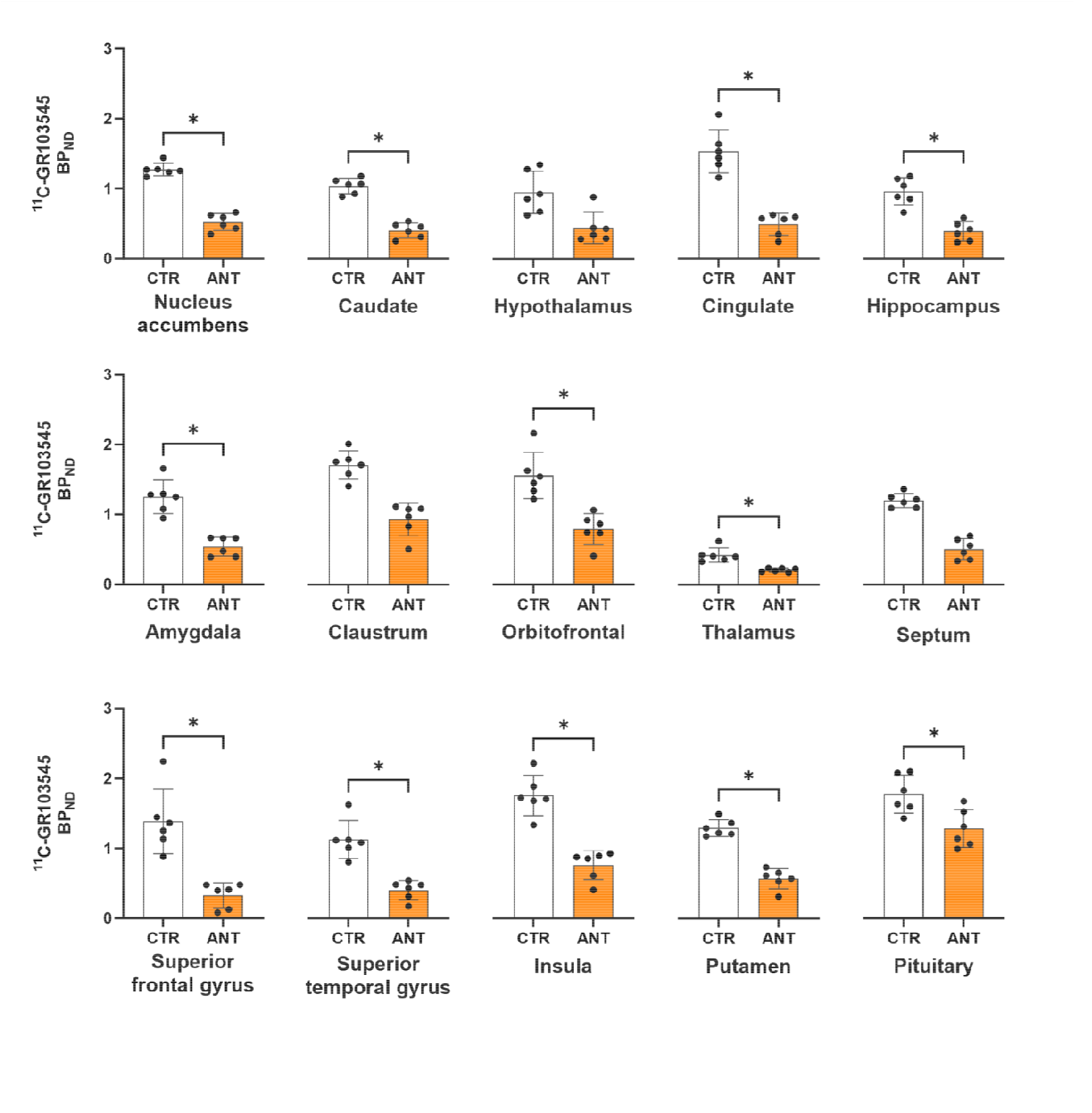
BP_ND_ across the 15 volumes of interest (VOIs) calculated with simplified reference tissue model (SRTM) for baseline (CTR) and CERC-501 pretreatment (ANT), for N=6 animals each. * indicates significant difference after FDR. The points indicate individual datapoints, while the bars indicate the 95% confidence interval

**Fig. 4.**
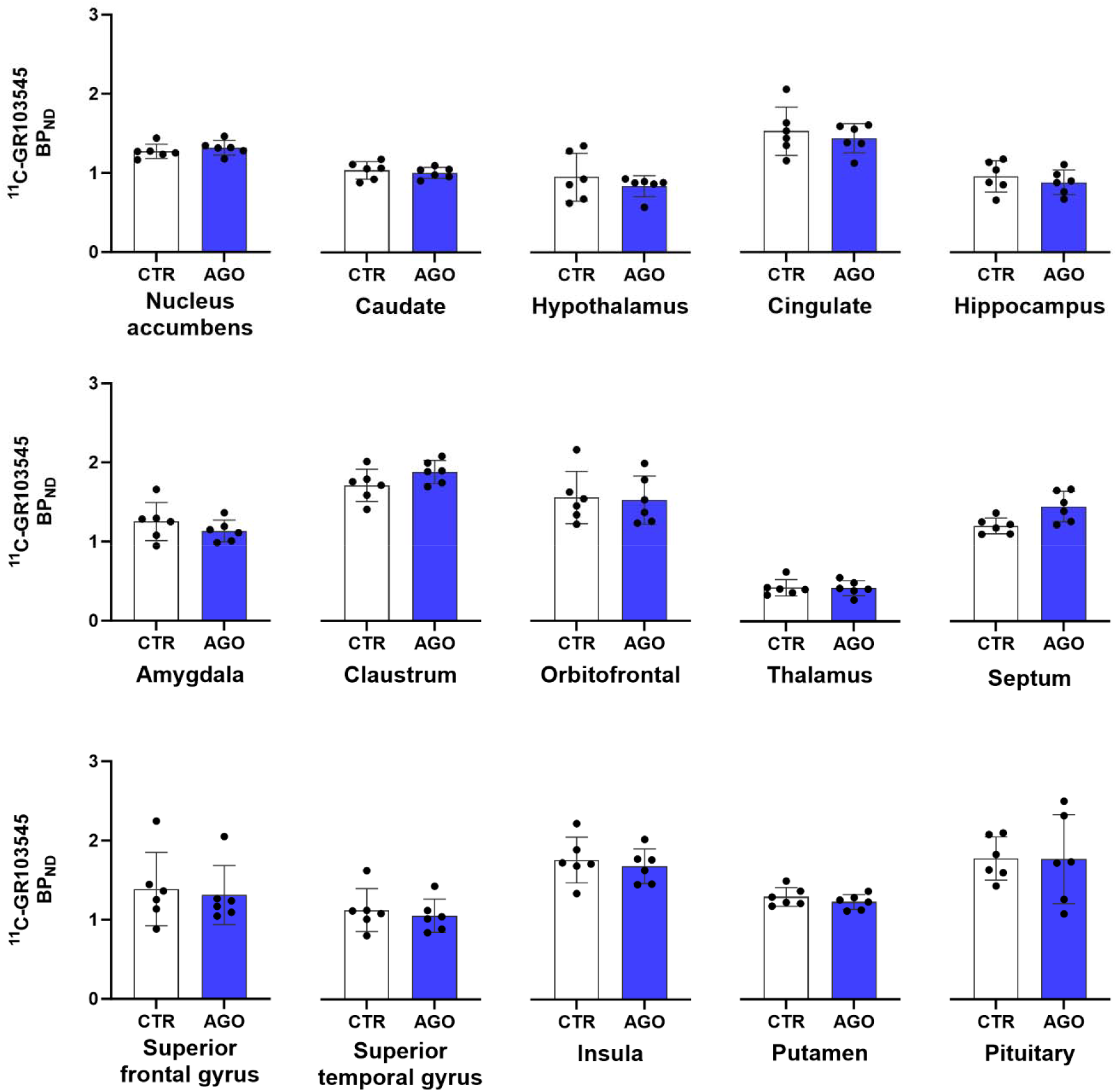
BP_ND_ across the 15 volumes of interest (VOIs) calculated with simplified reference tissue model (SRTM) for baseline (CTR) and U50,488 pretreatment (AGO), for N=6 animals each. While radiotracer binding differed in the different VOIs, there were no statistically significant differences between the CTR and AGO scans

### Pretreatment with antagonist CERC-501 significantly reduced binding across most VOIs when compared to baseline BP_ND_

There was a significant decrease in BP_ND_ post CERC-501 administration in comparison to baseline across most VOIs (average reduction of 55.99%; **Figure 2**). The superior frontal gyrus showed the largest reduction from baseline (76.25%, p=0.004) while the pituitary gland had the lowest reduction from baseline (27.69%, p=0.021) (**Supplementary Table S1**). Reductions in the claustrum, septum, and hypothalamus were not significant.

### Pretreatment with agonist U50,488 had no impact on ^11^C-GR103545 binding

KOR blockade by U50,488 appeared VOI-dependent on visual inspection, but was not significant across any analyzed VOI when compared to baseline (p>0.050). The overall reduction in BP_ND_ was 1.97% across the 15 VOIs with reductions ranging from 12.05% for the hypothalamus to 0.53% for the pituitary gland. Increased (but not significant) BP_ND_ was observed in the septum (−20.27%), claustrum (−9.99%) and nucleus accumbens (−3.48%) (**Supplementary Table S1**).

### High association for BP_ND_ between SRTM and LRTM

The LRTM results for the antagonist and agonist studies are provided in **Supplementary Section S2**. There was a strong positive correlation (R=0.95, p<0.001) between BP_ND_ assessed by SRTM versus the LRTM, with the latter showing a larger, but insignificant, overall reduction (**Supplementary Figures S2, S3, Supplementary Table S2**).

## Discussion

This study is the first to analyze KOR binding in titi monkeys using PET. The distribution of ^11^C-GR103545 in the titi monkey brain, as evaluated with BP_ND_, was consistent with KOR distribution from autoradiography [18]. Blocking of the KOR with antagonist CERC-501 significantly reduced ^11^C-GR103545 binding, supporting its role in KOR modulation. U50,488 pretreatment however did not impact ^11^C-GR103545 binding. The socially monogamous titi monkey demonstrates a unique combination of physiological similarity and complex social behaviors, analogous to humans, such as, elevated oxytocin (OT) levels upon separation from their partner [10, 19]. Given that the kappa opioid system (that modulates OT) may also contribute to the response to social separation, or “grief” there is interest in testing KOR interventions in this translatable model [8], thereby warranting the need for this current study.

The reduction in BP_ND_ after CERC-501 pretreatment was observed across all brain VOIs. These results are consistent with previously published ^11^C-GR103545 antagonist blocking experiments performed with naltrexone (an opioid antagonist) and PF04455242 (a KOR antagonist) in baboons [20] and humans [21], respectively. Self-blocking of ^11^C-GR103545 in rhesus monkeys [22], and blocking using a range of antagonists and agonists in rats [15], have also reported significant reductions in BP_ND_ and distribution volume (V_T_), respectively. The high selectivity of CERC-501 for mammalian KOR [23] has culminated in an ongoing phase 3 clinical trial for major depressive disorder (NCT06514742) and suggests that ^11^C-GR103545 BP_ND_ in titi monkeys indeed reflects KOR binding.

Conversely, U50,488 pretreatment resulted in no difference in BP_ND_ within the assessed brain VOIs. These results differ from the significant U50,488 blockade of KOR in rats [15]. There could be several reasons underlying this discrepancy. First, the KOR in titi monkeys has not yet been fully characterized structurally and functionally. Like humans [24] and macaques [25], titi monkeys may express multiple variants of the KOR [24] for which the agonist and radiotracer have differential binding. Second, it is plausible that species-level differences in receptor structure or post translational modifications may impact U50,488 binding affinity, causing reduced blockade [26]. Lastly, although our protocol for pretreatment was based on previous blocking studies [15, 20, 21], the timing, dose, or combination of U50,488 administration prior to radiotracer injection may have been suboptimal for the titi monkey. However, given the analgesic and aversive effects of U50,488 [27], a higher dose could not be justified given considerations of animal safety.

The use of reference tissue models for this analysis circumvents challenges associated with arterial blood sampling in a longitudinal experimental paradigm in a monkey model to study stress response. While the utility of the cerebellum as a reference region has been debated [12, 21, 22], kinetics justifying its use were reported in rats and baboons [15, 20].

Several radiotracers have been developed to evaluate KOR fluctuations *in vivo* with PET [28, 29, 30]. ^11^C-GR103545 was used in the present study due to its high binding affinity to KOR, as reported in rodent, non-human primate, and human studies [15, 20], [21, 22, 31], and the high correspondence between radiotracer uptake and KOR distribution identified *ex vivo* across species [20, 21, 22]. A comparison of the different radiotracers was outside the scope of our study but would be a future direction.

This study in titi monkeys using ^11^C-GR103545 was essential for establishing feasibility of utilizing this radiotracer in this translational animal model. The deployment of time course drug chase scans upon perturbing the KOR receptor with blocking agents was critical to analyze the impact on KOR activity. Lastly, this study lays the foundation for further experiments to analyze KOR binding dynamics longitudinally and in a post-intervention setting in the highly relevant titi monkey model of social attachment.

