## Supplemental data for "Mapping Kappa Opioid Receptor Binding in Titi Monkeys with ^11^C-GR103545 PET"

S1 **Methods**
   S1.1 **Animal upkeep and housing:**

All experiments with animals were approved by the UC Davis Institutional Animal Care and Use Committee and used facilities fully accredited by AAALAC International. The N=6 titi monkeys, born and housed at the California National Primate Research Center, were in cages (1.2 m × 1.2 m × 2.1 m) on a 12:12 light:dark cycle with temperature maintained at 21°C. All subjects were socially housed. Animals were fed a diet of monkey chow, banana, rice cereal, apple, and carrot twice a day.

S1.2 **Positron Emission Tomography (PET) radiotracer synthesis and acquisition**
S1.3 The PET radiotracer ^11^C-GR103545 was synthesized with a mean molar activity of 155.02±30.63 PBq/mol at the end of synthesis. Titi monkeys were anesthetized with 1-2% isoflurane and placed head-first on a custom bed of a dedicated brain PET scanner (PiPET, Brain Biosciences, Rockville, MD).

PET scans involved a 110-min dynamic emission data acquisition that began approximately 15 sec prior to injection of the radiotracer. Reconstructed data were generated with a framing of 6x10s, 8x30s, 5x60s, 4x300s, 8x600s and an isotropic voxel size of 0.8 mm^3^. A single-pass “cold” transmission scan was acquired for attenuation and scatter correction before the injection of the radiotracer.

S1.4 **Magnetic Resonance Imaging (MRI) acquisition**

T_1_-weighted anatomical MRs were acquired with a 1.5T GE Signa LX9.1 scanner (General Electric Corporation, Milwaukee, WI) and a 3” surface coil. The MR acquisition involved a 3D spoiled gradient echo (SPGR) pulse sequence in the coronal plane using the following parameters: echo time (TE)=7.9ms, repetition time (TR)=22.0ms, flip angle=30.0°, field of view=8cm, number of excitations=3, matrix=256 × 256, resulting in a pixel size of 0.3125 x 0.3125mm^2^ and a slice thickness of 1mm.

S1.5 **Image post processing and kinetic modeling:**

Each animal’s anatomical MR was brought to a titi monkey reference space by rigid registration. Anatomical MRs were then skull stripped using BrainSuite (v23a) and N4ITK bias-corrected. A titi monkey brain atlas was first aligned to each animal’s anatomical MR via rigid and affine transforms (mutual information-based cost function), and then warped via non-rigid, diffeomorphic deformations using the symmetric normalization (SyN) algorithm with cross correlation as the similarity metric and a B-spline based parameterization (Advanced Normalization Tools). The warped labels were further modified (where needed) by an expert in neuroanatomy. The PET data were motion-corrected via rigid registration to reference frame number 24. The motion-corrected PET was then manually rigid registered to its anatomical MRI on PMOD (v4.4, PMOD Technologies, Zürich, Switzerland). Kinetic modeling was then performed with simplified reference tissue model (SRTM), and Logan reference tissue model (LRTM) to obtain region-specific, non-displaceable binding potential (BP_ND_). **Figure S1** shows the image processing pipeline utilized in the study.

S1.6 **Logan reference tissue modeling**

Similar to SRTM, the cerebellum was used as the reference region. The average tissue-to-plasma clearance k_2_' was estimated from SRTM and the analysis was performed for the same volumes of interest (VOIs) as the main paper, namely, the nucleus accumbens, caudate, hypothalamus, cingulate, hippocampus, amygdala, claustrum, orbitofrontal cortex, thalamus, septum, superior frontal gyrus, superior temporal gyrus, insula putamen and pituitary gland.

S2 **Results**
   S2.1 **Pretreatment with antagonist CERC-501 significantly reduced binding across several VOIs when compared to baseline BP_ND_ as calculated with LRTM.**

Antagonist administration caused overall 59.68% reduction in BP_ND_. There was a significant decrease in binding when compared to baseline across all of the VOIs except the pituitary, claustrum and septum. Similar to SRTM, the superior frontal gyrus had the largest reduction from baseline (78.79%, p=0.021). Conversely, the pituitary had the lowest reduction from baseline (23.21%, p=0.149).

S2.2 **Pretreatment with agonist U50,488 had VOI specific changes in binding when compared to baseline BP_ND_ as calculated with LRTM.**

While an overall reduction of 6.03% was observed after agonist pretreatment, the changes were VOI dependent and not significant across any VOI. Conversely, 10 out of the 15 VOIs analyzed reported increased BP_ND_ when compared to baseline. The reductions ranged from the hypothalamus (12.05%, p=0.564) to the hippocampus (0.46%, p >0.999). Conversely, increased BP_ND_ ranged from the claustrum (-21.41%, p=0.083) to the thalamus (-0.16%, p=0.564).

S2.3 **BP_ND_ measured from LRTM correlated with that from the SRTM analysis.**

Given the strong positive correlation (R=0.95, p<0.001) between the two methods, notable differences include; no significant differences in the BP_ND_ of the pituitary gland and significant differences in the hypothalamus for CERC-501 pretreatment, with LRTM but not for SRTM. Lastly, both pretreatments had a larger overall reduction with the LRTM (antagonist: 59.68%, agonist: 6.03%) compared to SRTM (antagonist: 55.99%, agonist: 1.97%). However, only 5 out of the 15 VOIs had decreased BP_ND_ from agonist pretreatment, compared to the 12 from SRTM.

**Figures:**

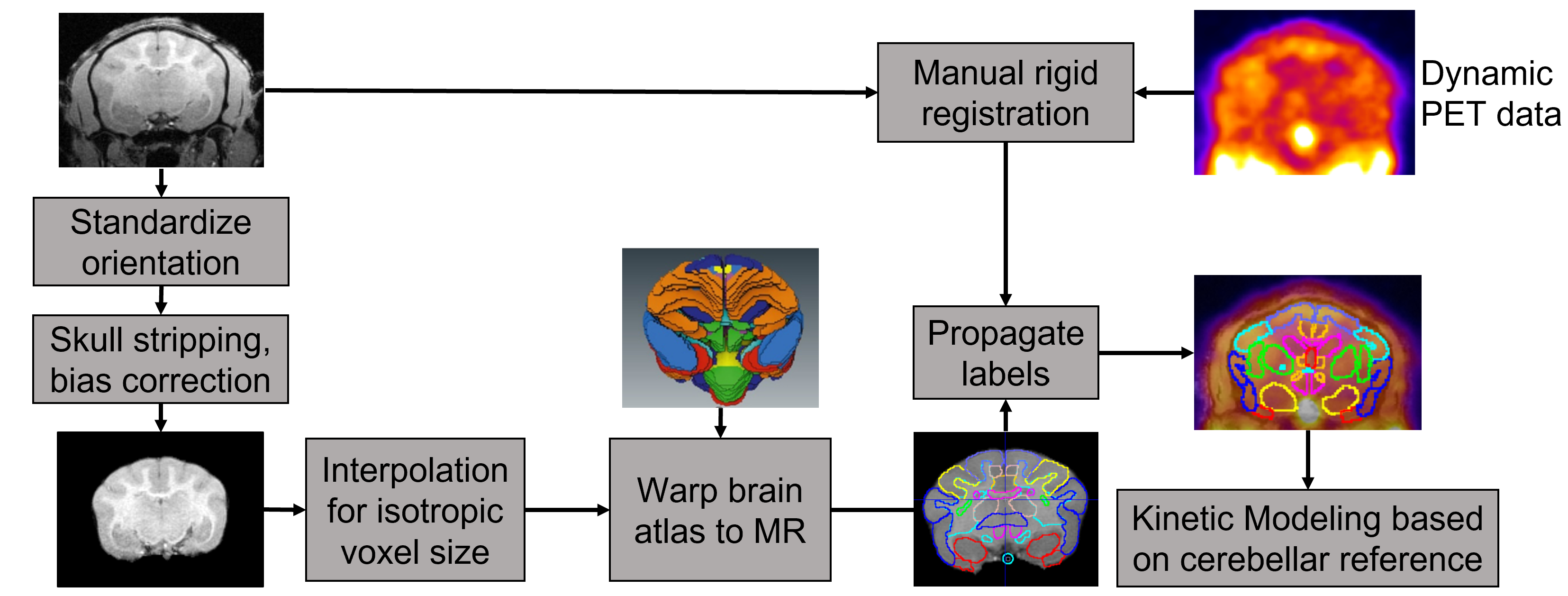

**Fig S1** Image post processing pipeline that involved warping of a titi monkey brain atlas to skull stripped, bias corrected anatomical MRs of each individual monkeys. Motion corrected PET was rigid registered to the anatomical MRI. Kinetic modeling was performed to calculate non-displaceable binding potential (BP_ND_) using simplified reference tissue model (SRTM), and Logan reference tissue model (LRTM), each using the cerebellum as the reference region. For LRTM, the average tissue-to-plasma clearance k_2_' was estimated from SRTM

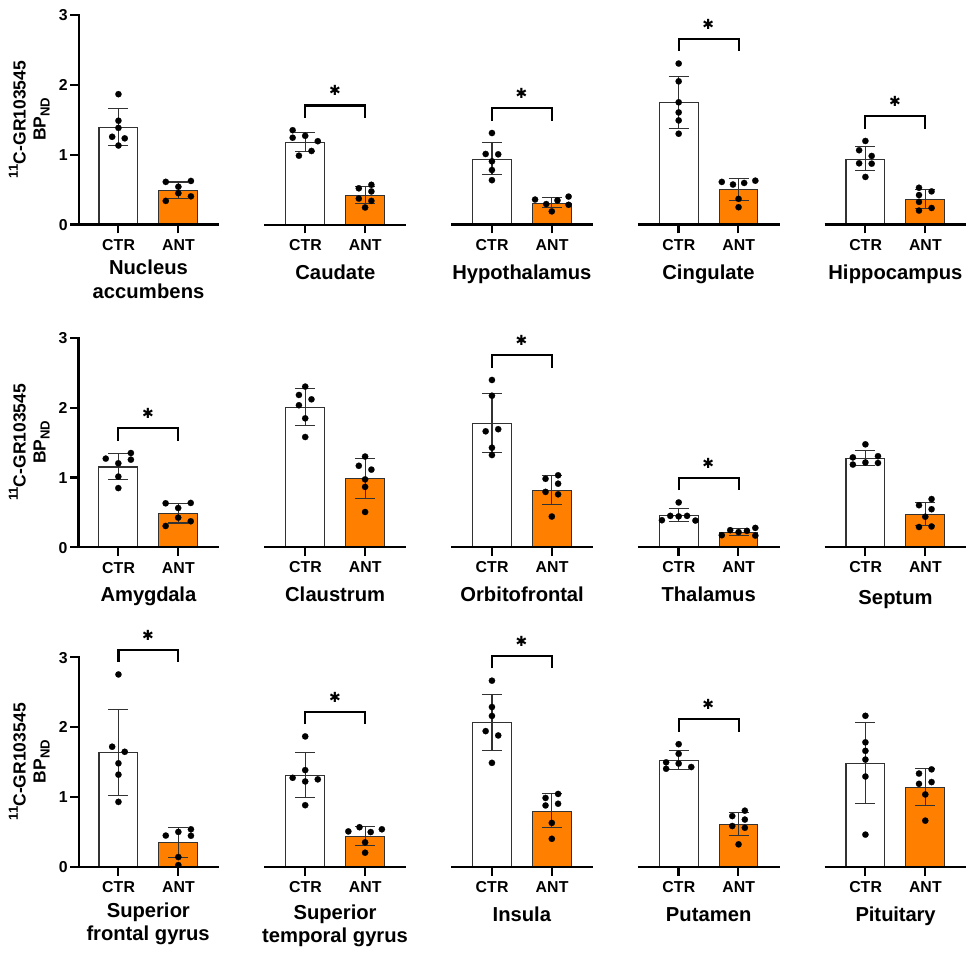

**Fig S2** BP_ND_ across the 15 volumes of interest (VOIs) calculated with Logan reference tissue model (LRTM) for baseline (CTR) vs antagonist pretreatment (ANT), for N=6 animals each.

* indicates significant difference across groups after FDR correction. The points indicate individual datapoints, while the bars indicate the 95% confidence interval

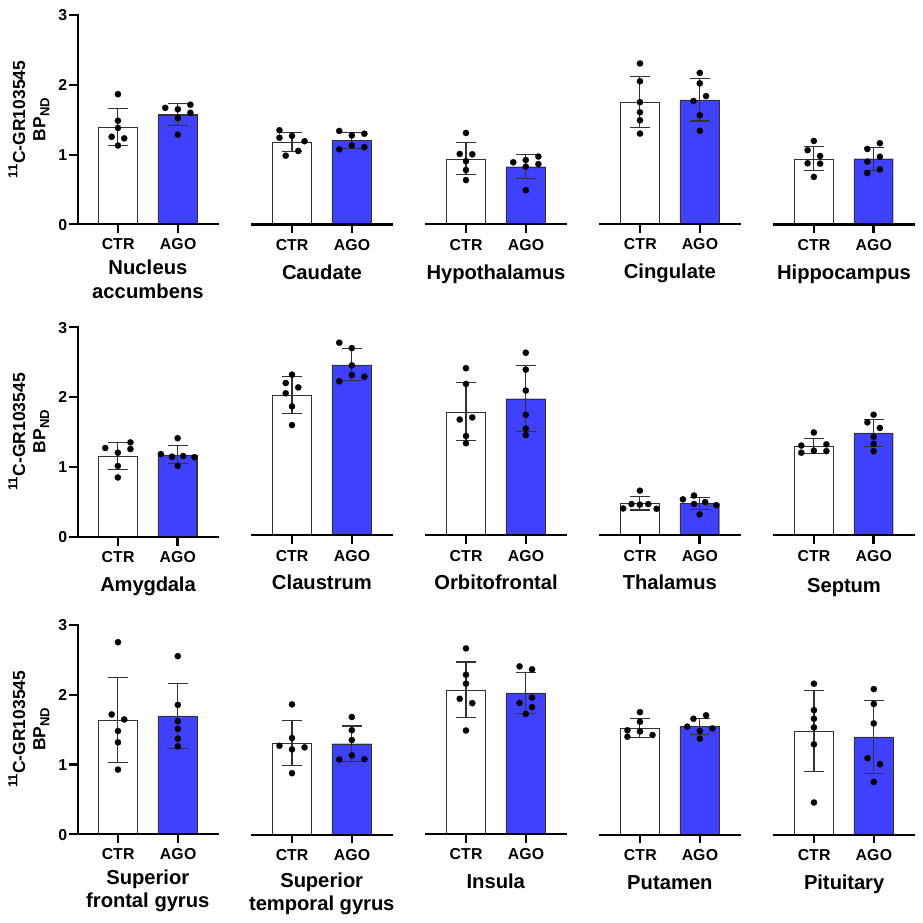

**Fig S3** BP_ND_ across the 15 volumes of interest (VOIs) calculated with Logan reference tissue model (LRTM) for baseline (CTR) vs agonist pretreatment (AGO), for N=6 animals each. While radiotracer binding differed in the different VOIs, there were no statistically significant differences between the CTR and AGO scans

**Tables:**

| **VOIs** | **Baseline** | **Antagonist** | | **Agonist** | |
| --- | --- | --- | --- | --- | --- |
|  | **Mean BP_ND_** | **Mean BP_ND_** | **p value (ANT vs CTR)** | **Mean BP_ND_** | **p value (AGO vs CTR)** |
| Amygdala | 1.25 | 0.54 | **0.004*** | 1.14 | 0.564 |
| Caudate | 1.04 | 0.41 | **0.004*** | 1.01 | 0.564 |
| Cingulate | 1.53 | 0.49 | **0.004*** | 1.44 | 0.564 |
| Claustrum | 1.72 | 0.94 | 0.083 | 1.89 | 0.083 |
| Hippocampus | 0.96 | 0.39 | **0.004*** | 0.89 | 0.564 |
| Hypothalamus | 0.95 | 0.45 | 0.043 | 0.84 | 0.564 |
| Insula | 1.76 | 0.76 | **0.004*** | 1.68 | 0.564 |
| Nucleus accumbens | 1.27 | 0.52 | **0.021*** | 1.32 | 0.564 |
| Orbitofrontal cortex | 1.56 | 0.79 | **0.004*** | 1.53 | 0.564 |
| Pituitary | 1.78 | 1.29 | **0.021*** | 1.77 | 0.564 |
| Putamen | 1.30 | 0.57 | **0.009*** | 1.23 | >0.999 |
| Septum | 1.21 | 0.51 | 0.083 | 1.45 | 0.083 |
| Superior frontal gyrus | 1.39 | 0.33 | **0.004*** | 1.31 | 0.564 |
| Superior temporal gyrus | 1.13 | 0.41 | **0.004*** | 1.06 | 0.564 |
| Thalamus | 0.43 | 0.21 | **0.021*** | 0.42 | 0.564 |

* indicates significance after FDR correction

**Table S1:** BP_ND_ calculated with simplified reference tissue model (SRTM) averaged across N=6 titi monkeys in the 15 volumes of interest (VOIs) scanned at baseline and after antagonist, CERC-501 as well as agonist U50,488 pretreatment, their corresponding p values upon comparing baseline BP_ND_ to BP_ND_ after antagonist/agonist pretreatment. The reported p-values are before FDR correction, while the asterisk denotes comparisons that survived FDR correction.

| **VOIs** | **Baseline** | **Antagonist** | | **Agonist** | |
| --- | --- | --- | --- | --- | --- |
|  | **Mean BP_ND_** | **Mean BP_ND_** | **p value (ANT vs CTR)** | **Mean BP_ND_** | **p value (AGO vs CTR)** |
| Amygdala | 1.16 | 0.49 | **0.004*** | 1.17 | 0.564 |
| Caudate | 1.19 | 0.42 | **0.021*** | 1.21 | 0.564 |
| Cingulate | 1.75 | 0.50 | **0.004*** | 1.78 | 0.564 |
| Claustrum | 2.01 | 0.99 | 0.083 | 2.44 | 0.083 |
| Hippocampus | 0.95 | 0.37 | **0.009*** | 0.94 | >0.999 |
| Hypothalamus | 0.94 | 0.31 | **0.004*** | 0.83 | 0.564 |
| Insula | 2.07 | 0.81 | **0.009*** | 2.03 | >0.999 |
| Nucleus accumbens | 1.39 | 0.49 | 0.043 | 1.58 | 0.248 |
| Orbitofrontal cortex | 1.78 | 0.82 | **0.009*** | 1.96 | >0.999 |
| Pituitary | 1.48 | 1.14 | 0.149 | 1.40 | 0.773 |
| Putamen | 1.53 | 0.61 | **0.009*** | 1.55 | >0.999 |
| Septum | 1.28 | 0.48 | 0.061 | 1.47 | 0.149 |
| Superior frontal gyrus | 1.64 | 0.35 | **0.021*** | 1.70 | 0.564 |
| Superior temporal gyrus | 1.31 | 0.44 | **0.004*** | 1.31 | 0.564 |
| Thalamus | 0.46 | 0.22 | **0.021*** | 0.46 | 0.564 |

* indicates significance after FDR correction

**Table S2:** BP_ND_ calculated using with Logan reference tissue model (LRTM) averaged across N=6 titi monkeys in the 15 volumes of interest (VOIs) scanned at baseline and after antagonist, CERC-501 as well as agonist U50,488 pretreatment, their corresponding p values upon comparing baseline BP_ND_ to BP_ND_ after antagonist/agonist pretreatment. The reported p-values are before FDR correction, while the asterisk denotes comparisons that survived FDR correction.
